# The complete mitochondrial genome of endangered Assam Roofed Turtle, *Pangshura sylhetensis* (Testudines: Geoemydidae): Genomic features and Phylogeny

**DOI:** 10.1101/827782

**Authors:** Shantanu Kundu, Vikas Kumar, Kaomud Tyagi, Kailash Chandra

## Abstract

Assam Roofed Turtle, *Pangshura sylhetensis* is an endangered and least studied species endemic to India and Bangladesh. The genomic feature of *P. sylhetensis* mitogenome is still anonymous to the scientific community. The present study decodes the first complete mitochondrial genome of *P. sylhetensis* (16,568 bp) by using next-generation sequencing. This *de novo* assembly encodes 13 Protein-coding genes (PCGs), 22 transfer RNAs (tRNAs), two ribosomal RNAs (rRNAs), and one control region (CR). Most of the genes were encoded on the majority strand, except NADH dehydrogenase subunit 6 (nad6) and eight tRNAs. Most of the PCGs were started with an ATG initiation codon, except for Cytochrome oxidase subunit 1 (cox1) and NADH dehydrogenase subunit 5 (nad5) with GTG. The study also found the typical cloverleaf secondary structure in most of the tRNA genes, except for serine (trnS1) with lack of conventional DHU arm and loop. Both, Bayesian and Maximum-likelihood topologies showed distinct clustering of all the Testudines species with their respective taxonomic ranks and congruent with the previous phylogenetic hypotheses (*Pangshura* and *Batagur* sister taxa). Nevertheless, the mitogenomic phylogeny with other amniotes corroborated the sister relationship of Testudines with Archosaurians (Birds and Crocodilians). Additionally, the mitochondrial Gene Order (GO) analysis indicated that, most of the Testudines species showed plesiomorphy with typical vertebrate GO.

## Introduction

The evolution of living organisms is a continuous process over generations and difficult to understand by measuring with a distinct hypothesis [1]. Several biological as well as environmental factors play an important role to change the descendent gene of a common ancestor that subsequently shifted into the next offspring’s or new species. Indeed, the genetic traits of a certain population drift randomly and evidenced to be gradually leads by the natural selection. Nevertheless, due to this high potency and tremendous success of genetic information, the evolutionary pattern of many living taxa are now well-understood through phylogenetic assessment. Hence, the world-wide collaborative effort of biologists ‘Tree of Life (ToL) project’ has been launched to illuminate the biodiversity and their evolutionary history [2].

Testudines (turtles, tortoises, and terrapins) are one of the oldest living organisms in the earth with an extended evolutionary history [3,4]. Besides the morphology, the genetic approach has been repeatedly applied to address the systematics of Testudines [5–7]. Both nuclear and mitochondrial genes have been extensively utilized to perceive the knowledge on Testudines phylogeny and genetic diversity [8–10]. Nonetheless, the Testudines mitogenomes were also scrutinized to understand their evolution and furnished the evidence of ‘Turtle-Archosaur affinity’ [11,12]. Additionally, to address the evolution of turtles within amniotes and strengthen the Vertebrate tree of life, the phylogenomic approach was also evaluated [13,14]. However, the phylogeny of Testudines is yet to be reconciled by mining of large-scale molecular data to know their precise evolutionary scenario.

Testudines mitochondrial genomes are usually circular and double stranded with 16-19 kb in size contains 37 genes [15–19]. Remarkably, some of the reptile species including turtles, have variable numbers of genes in their mitogenome due to the losing of protein-coding genes (PCGs), duplication of control regions (CRs), and occurrence of multiple contiguous numbers of one or more transfer RNA genes (tRNAs) [20]. Besides the structural features of different genes, the gene orders (GOs) in mitogenome have also evidenced to be the signature of defining clades at different taxonomic levels in both vertebrates and invertebrates [21–25]. In general, the arrangements of GO are defined by transposition, inversion, inverse transposition, and tandem duplication and random loss (TDRL) mechanisms in comparisons with the typical ancestral GO [26]. Moreover, these evolutionary events also help to define the plesiomorphic/apomorphic status and transformational pathways of mitogenome [27,28]. Thus, apart from the conventional analysis; to infer the evolutionary pathways leading to the detected diversity of GOs, it is essential to test phylogeny and GO conjointly [29,30]. As of now, more than 100 Testudines species mitogenomes of 56 genera within 12 families and two sub-orders were generated worldwide to address several phylogenetic questions. However, the combined study on GO and phylogeny has never been assessed for Testudines to discuss the evolutionary tracts.

The evolutionarily distinct genus *Pangshura* is known by four extant species from Southeast Asian countries [31]. Among them, the Assam Roofed turtle, *Pangshura sylhetensis* is a highly threatened species and categorized as ‘Endangered’ in the International Union for Conservation of Nature (IUCN) Red data list [32]. The distribution of *P. sylhetensis* is restricted to Bangladesh and India [33,34]. Although, the partial segments of mitochondrial genetic information are publicly available, but the mitogenomic information on this species is still anonymous to the scientific communities. Therefore, the present study aimed to generate the complete mitochondrial genome of *P. sylhetensis* and execute the comparative analysis with other Testudines belonging to both sub-orders (Cryptodira and Pleurodira). Here, we used phylogenetic (Bayesian and Maximum Likelihood) and GO analyses for better insights into the Testudines evolutionary scenario.

## Materials and Methods

### Ethics statement and sample collection

To conduct the field survey and biological sampling, prior permission was acquired from the wildlife authority of Arunachal Pradesh (Letter No. SFRI/APBB/09-2011-1221-1228 dated 22.07.2016) and Zoological Survey of India, Kolkata (Letter No. ZSI/MSD/CDT/2016-17 dated 29.07.2016). No turtle specimens were sacrificed in the present study. The *P. sylhetensis* specimen was collected from the tributaries of Brahmaputra River (Latitude 27.50 N and Longitude 96.24 E) from the nearby localities of Namdapha National Park in northeast India (Fig. 1). The species photograph was taken by the first authors (S.K.) and edited manually in Adobe Photoshop CS 8.0. The country level topology map has been downloaded from DIVA-GIS Spatial data platform (http://www.diva-gis.org/datadown) and overlaying by ArcGIS 10.6 software (ESRI®, CA, USA). The range distribution was edited manually in Adobe Photoshop CS 8.0 followed by the published record [35]. The blood sample was collected in sterile condition from the hind limb of the live individual by using micro-syringe and stored in EDTA vial at 4°C. Subsequently the specimen was released back in the same eco-system with ample care and attention.

**Fig 1.**
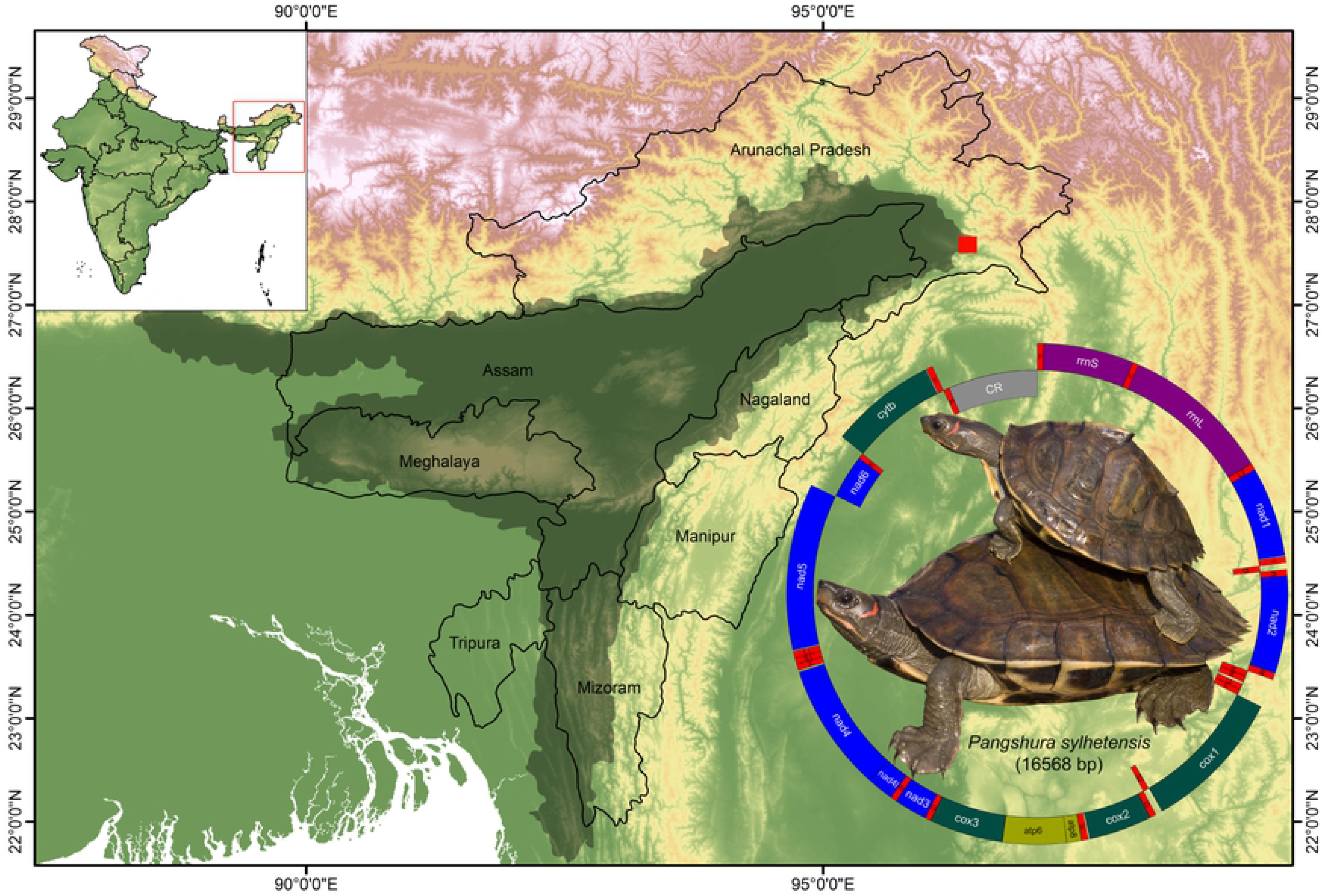
The spatial range distribution (marked by gray shadow) and collection locality (27.50 N 96.24 E marked by red square) of *P. sylhetensis*. The species photograph and mitochondrial genome of *P. sylhetensis*. Protein-coding genes are marked by blue, deep green, and yellowish-green color boxes, rRNA genes are marked by violet color boxes, tRNA genes are marked by red color boxes, and control region is marked by gray color box.

### Mitochondrial DNA extraction and sequencing

The collected blood sample (20 μl) was thoroughly blended with 1 ml buffer (0.32 M Sucrose, 1 mM EDTA, 10 mM TrisHCl) and centrifuged at 700 × g for 5 min at 4°C to remove nuclei and cell debris. The supernatant was collected in 1.5 ml eppendorf tubes and centrifuged at 12,000 × g for 10 min at 4°C to precipitate the mitochondrial pellet. The pellet was re-suspended in 200 μl of buffer (50 mM TrisHCl, 25 mM of EDTA, 150 mM NaCl), with the addition of 20 μl of proteinase K (20 mg/ml) followed by incubation at 56°C for 1-2 hr and the mitochondrial DNA was extracted by Qiagen DNeasy Blood & Tissue Kit (QIAGEN Inc.). The DNA quality was visualized in 1% agarose gel electrophoresis, and the concentration was quantified by NANODROP 2000 spectrophotometer (Thermo Scientific).

### Mitogenome assembly and annotation

Complete mitochondrial genome sequencing and *de novo* assembly was carried out at Xcelris Labs Limited, Gujarat, India (http://www.xcelrisgenomics.com/). The genome library was sequenced using the Illumina platform (2×150bp PE chemistry) with the average size of library was 545bp, to generate ~3 GB data (Illumina, Inc, USA). The total yield of the Number of Paired end was 26,263,128 with the maximum data of 3.78 GB. The raw reads were handled using cutadapt tool (http://code.google.com/p/cutadapt/) for adapters and low quality bases trimming with cutoff of Phread quality score (Q score) of 20. The high quality reads were down sampled to 2 million reads using Seqtk program (https://github.com/lh3/seqtk). High quality paired end data was assembled with NOVOPlasty v2.6.7 with default parameters [36]. The mitogenome of *Mauremys reevesii* (accession no. KJ700438) was used as a reference seed sequence for *de novo* assembly with 16568 bp (~16.5Kb) single contig. To validate the assembly obtained from NOVOPlasty assembler, similarity search was carried out in GenBank database using BLASTn v2.2.28 search (https://blast.ncbi.nlm.nih.gov) algorithm. Further, the contig was subjected to confirm by MITOS v806 online webserver (http://mitos.bioinf.uni-leipzig.de). The DNA sequences of the protein coding genes (PCGs) were primarily translated into the putative amino acid sequences on the basis of the vertebrate mitochondrial genetic code. The exact initiation and termination codons were identified in ClustalX using other publicly available reference sequences of Testudines [37]. The de-novo mitogenome was submitted to the GenBank database through the Sequin submission tool (Fig. S1).

### Data set construction and comparative analysis

The spherical representation of the generated mitogenome of *P. sylhetensis* was plotted by CGView Server (http://stothard.afns.ualberta.ca/cgview_server/) with default parameters [38]. The strand direction and arrangements of each PCG, tRNA, and ribosomal RNA (rRNA) were also checked through MITOS online server. The overlapping regions and intergenic spacers between the neighbor genes were counted manually through Microsoft Excel. The tRNA genes of *P. sylhetensis* were verified in MITOS online server, tRNAscan-SE Search Server 2.0 (http://lowelab.ucsc.edu/tRNAscan-SE/) and ARWEN 1.2 [39,40]. The base composition of all stems (DHU, acceptor, TѱC, anticodon) were examined manually to discriminate the Watson-Crick, wobble, and uneven base pairing. On the basis of homology search in the Refseq database (https://www.ncbi.nlm.nih.gov/refseq/), 51 Testudines mitogenomes representing nine families of Cryptodira and three families of Pleurodira were downloaded from GenBank and incorporated in the dataset for comparative analysis (Table S1). The genome size and comparative analysis of nucleotide composition with other Testudines were calculated using MEGA6.0 [41]. The base composition skew was calculated as defined previously: AT skew = (A – T)/(A + T), GC skew = (G – C)/(G + C) [42]. The start and stop codons of PCGs were checked through the Open Reading Frame Finder web tool (https://www.ncbi.nlm.nih.gov/orffinder/). To determine the location for replication of the L-strand and putative secondary structures, all Testudines CRs were analyzed through online Mfold web server (http://unafold.rna.albany.edu). Due to the lack of proper annotation, the CRs of two species (*Amyda cartilaginea* and *Phrynops hilarii*) were missing and thus not incorporated in the comparative analysis.

### Phylogenetic and Gene Order (GO) analysis

To assess the Bayesian analysis (BA) and maximum-likelihood (ML) phylogenetic relationship, two datasets were prepared in the present analysis. The first dataset includes all the PCGs of 52 Testudines mitogenomes including *P. sylhetensis* (Table S1). As the targeted species is a Cryptodiran species, all the Pleurodiran species were treated as out-group taxa in the first dataset. Further, for reassess the existing hypothesis of Testudines phylogenetic position with other amniotes, all the PCGs of 11 database sequences (Whale: KU891394, Human: AP008580, Platypus: NC_000891, Opossum: AJ508398, Iguana: NC_002793, Lizards: AB080237, Snake: HM581978, Tuatara: AF534390, Bird: AP003580, Alligator: NC_001922, and Crocodile: HM488007) were acquired from the GenBank database to construct the second dataset (Table S1). Among them, the database sequences of aquatic Amniota (*Physeter catodon*: KU891394) was used as out-group taxa. The PCGs were aligned individually by codons using MAFFT algorithm in TranslatorX with L-INS-i strategy with GBlocks parameters [43]. Finally, the two datasets of all PCGs was concatenated using SequenceMatrix v1.7.84537 [44]. The suitable models for phylogenetic analysis were estimated by PartitionFinder 2 [45] at CIPRES Science Gateway V. 3.3 [46] (Table S2). The BA tree was built through Mr. Bayes 3.1.2 and the metropolis-coupled Markov Chain Monte Carlo (MCMC) was run for 100,000,000 generations with sampling at every 100^th^ generation and 25% of samples were discarded as burn-in [47]. The ML tree was constructed using IQ-Tree web server with 1000 bootstrap support [48]. Both BA and ML topologies for two datasets were further refined in iTOL v4. (https://itol.embl.de/login.cgi) tool for better illustration [49] and edited with Adobe Photoshop CS 8.0.

Further, to check the gene arrangement scenario, most contemporary TreeREx analysis was adopted to infer the evolutionary pathways within the Testudines, leading to the observed diversity of the GOs. TreeREx can easily distinguish the putative GOs at the internal nodes of a reference tree as it works in a bottom-up manner through the iterative analysis of triplets or quadruplets of GOs to decide all the GOs in the entire tree [30]. In the TreeREx analysis, the consistent nodes are considered to be most reliable and marked by green color, whereas the fallback nodes have the highest level of uncertainty marked by red color. In doing so, we used the default settings of TreeREx suggested on the website (http://pacosy.informatik.uni-leipzig.de/185-0-TreeREx.html) to analyze every node of the reference phylogenetic tree was defined by a GO. To finalize the gene arrangements dataset of 52 Testudines for TreeREx analysis (Table S3); the insertion, deletion, and duplication of genes within the database available mitogenomes were reminisced as discussed in the previous studies [50–53].

## Results and Discussion

### Mitogenome structure and organization

The mitogenome (16,568 bp) of endangered Assam Roofed turtle, *P. sylhetensis* was determined in the present study (GenBank accession no. MK580979). The mitogenome was encoded by 37 genes, comprising 13 PCGs, 22 tRNAs, two rRNAs, and a major non-coding CR. Among them, nine genes (NADH dehydrogenase subunit 6 and eight tRNAs) were located on the minority strand, while the remaining 28 genes were located on the majority strand (Table 1 and Fig. 1). In the present dataset, the length of mitogenome was varied from 15,339 bp (*A. cartilaginea*) to 19,403 bp (*Stigmochelys pardalis*). Out of 52 Testudines species, 43 species were showed strands symmetry as observed in typical vertebrates [54]. The gene arrangements scenarios within the Testudines species are discussed elaborately in the forthcoming subsection. The nucleotide composition of *P. sylhetensis* mitogenome was A+T biased (59.27%) as represented in other Testudines mitogenomes ranging from 57.76% (Pleurodiran species *P. hilarii*) to 64.19% (Cryptodiran species *Kinosternon leucostomum*) (Table S4). The A+T composition of *P. sylhetensis* PCGs was 58.77%. The AT skew and GC skew was 0.124 and −0.334 in the mitogenome of *P. sylhetensis*. The comparative analysis showed that the AT skew ranged from 0.087 (*Testudo graeca*) to 0.208 (*Carettochelys insculpta*) and GC skew from −0.296 (*S. pardalis*) to −0.412 (*Trionyx triunguis*) (Table S4). A total of 15 overlapping regions with a total length of 73 bp were identified in *P. sylhetensis* mitogenome. The longest overlapping region (21 bp) was observed between tRNA-Leucine (trnL1) and NADH dehydrogenase subunit 5 (nad5). Further, a total of 12 intergenic spacer regions with a total length of 129 bp were observed in *P. sylhetensis* mitogenome with a longest region (66 bp) between tRNA-Proline (trnP) and CR (Table 1).

**Table 1.** List of annotated mitochondrial genes of *Pangshura sylhetensis*.

| Gene | Direction | Location | Size (bp) | Anti-codon | Start codon | Stop codon | Intergenic Nucleotides |
| --- | --- | --- | --- | --- | --- | --- | --- |
| trnF | + | 1-69 | 69 | GAA | . | . | 0 |
| rrnS | + | 70-1031 | 962 | . | . | . | 0 |
| trnV | + | 1032-1100 | 69 | TAC | . | . | 11 |
| rrnL | + | 1112-2697 | 1586 | . | . | . | 1 |
| trnL2 | + | 2699-2774 | 76 | TAA | . | . | 0 |
| nad1 | + | 2775-3743 | 969 | . | ATG | TAG | -1 |
| trnI | + | 3743-3813 | 71 | GAT | . | . | -1 |
| trnQ | - | 3813-3883 | 71 | TTG | . | . | -1 |
| trnM | + | 3883-3951 | 69 | CAT | . | . | 0 |
| nad2 | + | 3952-4992 | 1041 | . | ATG | TAG | -2 |
| trnW | + | 4991-5064 | 74 | TCA | . | . | -1 |
| trnA | - | 5064-5132 | 69 | TGC | . | . | 1 |
| trnN | - | 5134-5207 | 74 | GTT | . | . | 27 |
| trnC | - | 5235-5300 | 66 | GCA | . | . | 0 |
| trnY | - | 5301-5371 | 71 | GTA | . | . | 1 |
| cox1 | + | 5373-6923 | 1551 | . | GTG | AGG | -12 |
| trnS2 | - | 6912-6982 | 71 | TGA | . | . | 0 |
| trnD | + | 6983-7052 | 70 | GTC | . | . | 0 |
| cox2 | + | 7053-7739 | 687 | . | ATG | TAG | 1 |
| trnK | + | 7741-7813 | 73 | TTT | . | . | 1 |
| atp8 | + | 7815-7982 | 168 | . | ATG | TAA | -10 |
| atp6 | + | 7973-8656 | 684 | . | ATG | TAA | -1 |
| cox3 | + | 8656-9440 | 785 | . | ATG | TA(A) | -1 |
| trnG | + | 9440-9507 | 68 | TCC | . | . | 1 |
| nad3 | + | 9509-9857 | 349 | . | ATG | GAA | 1 |
| trnR | + | 9859-9927 | 69 | TCG | . | . | 0 |
| nad4l | + | 9928-10224 | 297 | . | ATG | TAA | -7 |
| nad4 | + | 10218-11594 | 1377 | . | ATG | TAA | 14 |
| trnH | + | 11609-11677 | 69 | GTG | . | . | 0 |
| trnS1 | + | 11678-11744 | 67 | GCT | . | . | -1 |
| trnL1 | + | 11744-11816 | 73 | TAG | . | . | -21 |
| nad5 | + | 11796-13628 | 1833 | . | GTG | TAG | -8 |
| nad6 | - | 13621-14145 | 525 | . | ATG | AGG | -3 |
| trnE | - | 14143-14210 | 68 | TTC | . | . | 4 |
| cytb | + | 14215-15361 | 1147 | . | ATG | T(AA) | -3 |
| trnT | + | 15359-15430 | 72 | TGT | . | . | 0 |
| trnP | - | 15431-15501 | 71 | TGG | . | . | 66 |
| D-loop | . | 15568-16389 | 822 | . | . | . | . |

### Protein-coding genes

The total length of PCGs was 11,268 bp in *P. sylhetensis*, which represent 68.01% of complete mitogenome. The AT skew and GC skew was 0.056 and −0.345 in PCGs of *P. sylhetensis* (Table S4). Most of the PCGs of *P. sylhetensis* started with an ATG initiation codon; however the GTG initiation codon was observed in Cytochrome oxidase subunit 1 (cox1) and nad5. The TAG termination codon was used by four PCGs, TAA by four PCGs, AGG by two PCGs, and GAA by one PCG. The incomplete termination codons TA(A) and T(AA) were detected in Cytochrome oxidase subunit 3 (cox3) and Cytochrome b (cytb) gene respectively. The comparative analysis with other Testudines species revealed that the high frequency of initiation codons (ATN, GTG) were observed in NADH dehydrogenase subunit 1 (nad1), NADH dehydrogenase subunit 2 (nad2), NADH dehydrogenase subunit 3 (nad3), NADH dehydrogenase subunit 4L (nad4L), NADH dehydrogenase subunit 4 (nad4), nad5, and cytb. The frequency of ATG initiation codon is always higher in most of the PCGs (64.71% to 96.08%), except for cox1 with GTG start codon (72.55%). The higher frequency of complete termination codon was observed in eight PCGs, except five PCGs with incomplete termination codon (Table S5 and Fig. 2).

**Fig. 2.**
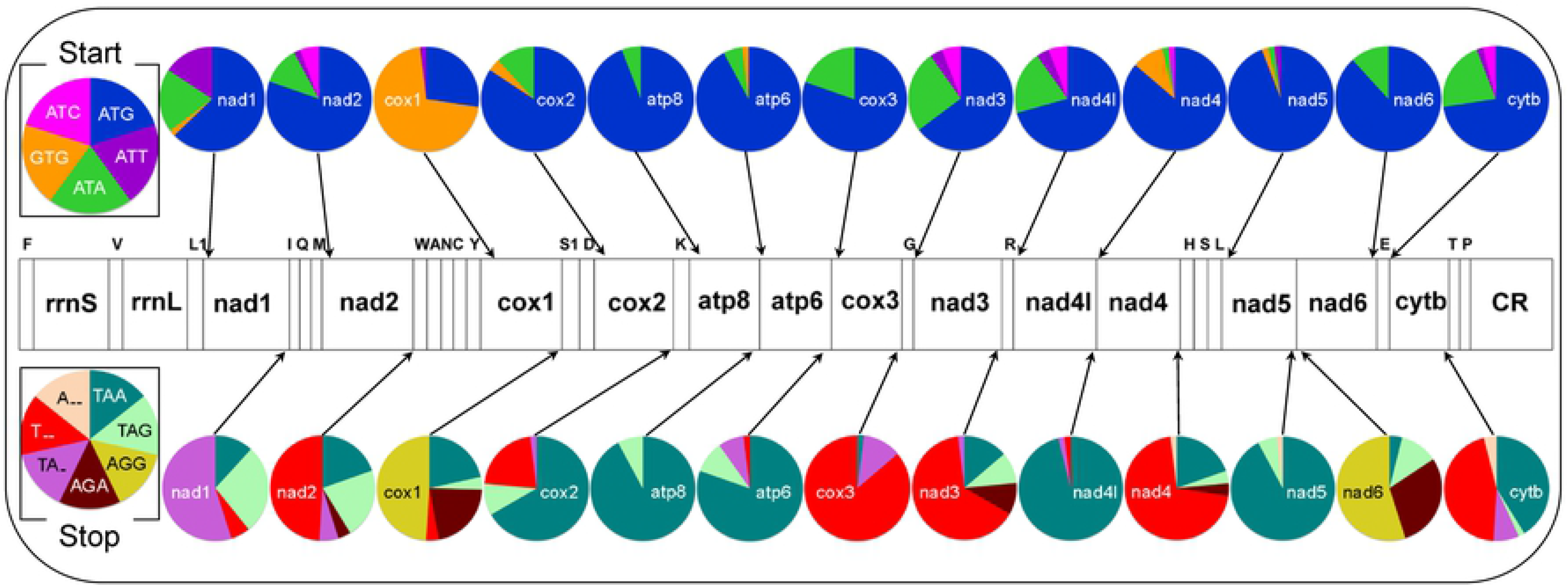
Frequency of start and stop codons for 13 protein-coding genes in the 52 Testudines mitogenomes.

### Ribosomal RNA and transfer RNA genes

The total length of two rRNA genes of *P. sylhetensis* was 2,560 bp with ranged from 1,611 bp (*Caretta caretta*) to 2,685 bp (*Pelodiscus sinensis*) in the present dataset. The AT content within rRNA genes was 58.98% with AT and GC skew was 0.272 and −0.173 respectively (Table S4). Total 22 tRNAs were found in the *P. sylhetensis* mitogenome with the total length of 1,550 bp. In other Testudines, the length of tRNAs was varied from 1,407 bp (*A. cartilaginea*) to 1,684 bp (*Platysternon megacephalum*). The AT content within tRNA genes was 59.87% with AT and GC skew was 0.017 and 0.057 respectively (Table S4). Among all the tRNA genes, 14 were present in majority strand and eight tRNA genes (trnQ, trnA, trnN, trnC, trnY, trnS2, trnE, and trnP) on minority strand. The anticodons of all the tRNA genes were also detected in the present study. Most of the tRNA genes were folded into classical cloverleaf structure except trnS1 (without Dihydrouridine (DHU) stem and loop (Fig. S2). The conventional base pairings (A=T and G≡C) were observed in most of the tRNAs [55]; however the wobble base pairing were observed in the stem of 13 tRNAs (trnL2, trnN, trnA, trnW, trnP, trnE, trnR, trnG, trnC, trnY, trnS2, trnK, trnQ) (Fig. S2).

### Control regions

The CR of *P. sylhetensis* was typically distributed with three functional domains, the termination associated sequence (TAS), the central conserved (CD), and the conserved sequence block (CSB) as observed in other vertebrate CRs [56]. As compared to the TAS and CSB domain with varying numbers of tandem repeats, the CD domain consisting with highly conserved sequences. Hence, the pattern of CR was varied among different vertebrates, including Testudines [56,57]. The total length of *P. sylhetensis* CR was 1,067 bp, ranging from 600 bp (*Lepidochelys olivacea*) to 3,885 bp (*S. pardalis*) in the present dataset. In *P. sylhetensis* CR, the AT and GC skew was −0.025 and −0.249 (Table S4). The TAS domain was ‘TACATA’, while the CSB domain was further divided into four regions: CSB-F (AGAGATAAGCAAC), CSB-1 (GACATA), CSB-2 (TTAAACCCCCCTACCCCCC), and CSB-3 (TCGTCAAACCCCTAAATCC). The CR is also acknowledged for the initiation of replication, and is positioned between trnP and trnF for most of the Testudines except *P. megacephalum* [50,52]. In, *P. sylhetensis* 27 bp nucleotides are present between CSB-1 and stem-loop structure, however in other species, it was ranging from zero to 90 bp (*T. triunguis*) (Fig. S3 and S4). Overall, the structural features of the replication of the L-strand and putative secondary structures are different in all Testudines species, which can be used as a species-specific marker.

### Phylogeny of Testudines mitogenomes

The phylogenetic position of Testudines in the Vertebrate tree of life has been evaluated in last few decades. Several approaches have been aimed to reconcile the phylogeny for better understanding of their origin and diversification [4]. Besides, morphological parameters the gene based topology have been widely measured for their identification, species delimitation, and population genetics assessment [58,59]. Nevertheless, to interpret the evolutionary scenario, the tested dataset should comprise with comprehensive genetic information on various taxa representing all extant lineages. Both morphological and molecular data corroborates to erect all the *Pangshura* species from the closest congeners of *Batagur* in earlier studies [31,60,61]. The mitogenomic data have successfully evidenced the phylogenetic relationships of many Testudines species, including the studied genus *Pangshura* under Geoemydidae family [62]. Till date, majority of species under sub-family Geoemydinae has been targeted throughout the world to generate their mitogenomes. Nevertheless, only two species mitogenomes of sub-family Batagurinae are presently available in the global database. Hence, the present study adds the *de novo* assembly third Batagurinae species (*P. sylhetensis*) in the global database. Both BA and ML phylogenies effectively discriminated all the studied Testudines species compiled in the first dataset with high bootstrap support and posterior probability (Fig. 3 and S5). The present phylogeny evidenced the sister relationship of *P. sylhetensis* with *Batagur trivittata* species as described in the previous phylogeny [62]. The other representative species are also showed close clustering within their respective families and sub-orders as well as consistent with the previous phylogenetic hypothesis [59,61]. The present mitogenomic phylogeny with concatenated 13 PCGs was adequate to generate a robust phylogeny and illuminate the relationship between *P. sylhetensis* with other Testudines species. More taxa from different taxonomic lineages from diverse distribution localities and their large-scale genetic data will be worthy to settle more exhaustive phylogeny and evolutionary connection.

**Fig 3.**
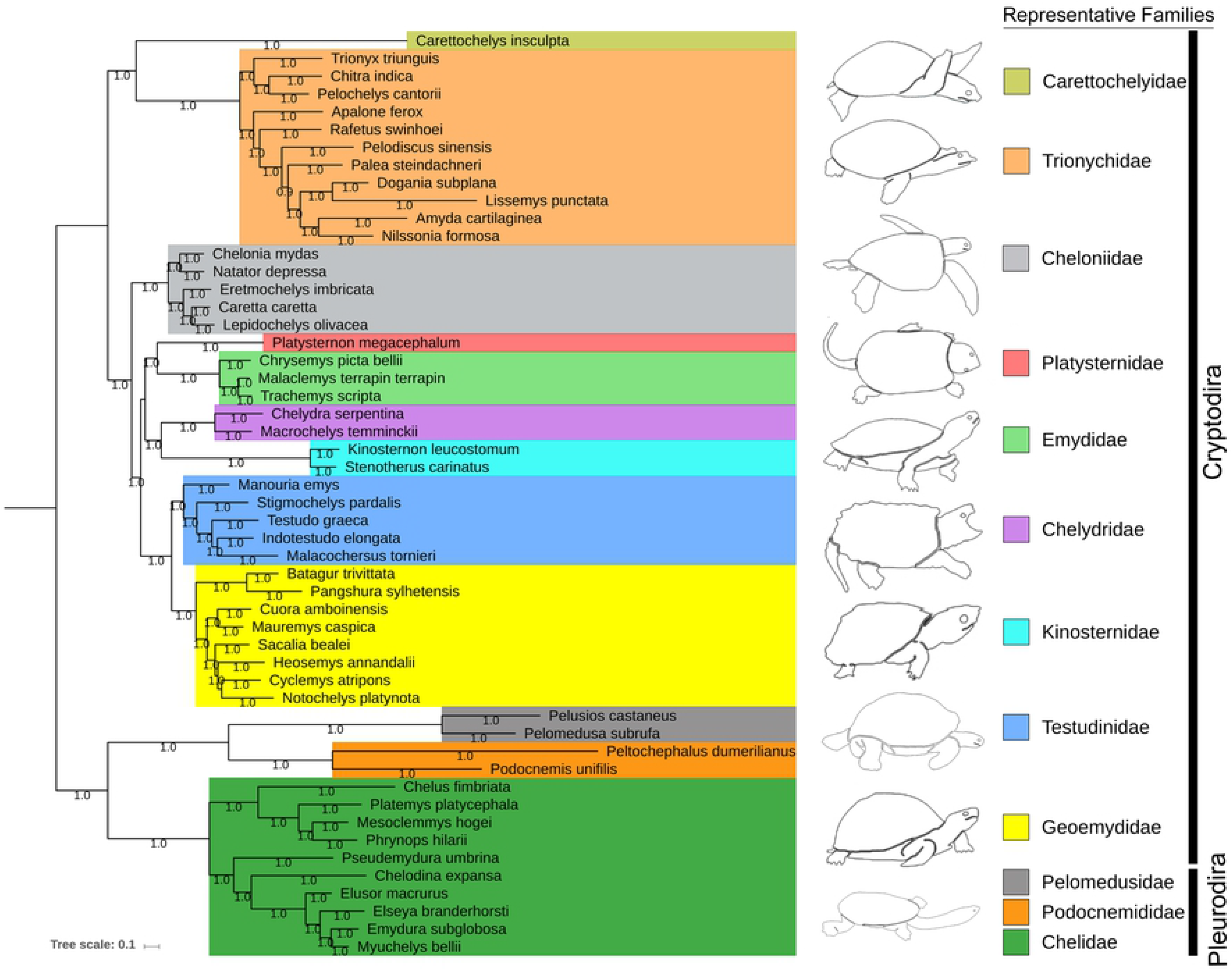
Bayesian (BA) phylogenetic tree based on the concatenated nucleotide sequences of 13 PCGs of 52 Testudines species showing the evolutionary relationship of *P. sylhetensis*. Color boxes indicate the family level clustering for the studied species. The BA tree is drawn to scale with posterior probability support values were indicated along with the branches. Representative organism line diagrams are acquired from cyberspace.

In addition, the evolutionary position of Testudines is still controversial in relationship with other amniotes [63–65]. To resolve the impediments on the placement of Testudines either in anapsids or diapsids, the mitogenome of Pleurodiran species has been analyzed previously and reject the placement of Testudines as most basal position in the amniote tree of life [11,66]. The mitogenome sequences have also been studied to evaluate the relationships between Archosaurians (birds and crocodilians) and Lepidosaurians (tuatara, snakes, and lizards) [67]. In recent past, the reptilian transcriptome based phylogenetic analysis also suggests that, Testudines are not the basal of extant reptiles but showed affinity towards other Archosaurians [68]. Further, the candidate nuclear protein-coding locus (NPCL) markers were also evaluated and demonstrated that, turtles are robustly recovered as the sister group of Archosauria (birds and crocodilians) and the evolutionary timescales was congruent with the Timetree of Life [69]. Later on, the phylogenomic approaches and ultra-conserved elements (UCEs) based analysis were endeavored to test the prevailing hypothesis and supports the sister relationships of Testudines with Archosaurians [13,70,71]. Hence, to affirm the existing phylogenetic hypothesis of Testudines with other amniotes, we constructed the phylogenetic tree of mitogenomes dataset (Testudines + other amniotes). Both BA and ML phylogeny of this dataset showed close clustering with birds, alligator, and crocodile in comparison with other amniotes with high bootstrap support and posterior probability (Fig. 4 and S6). Hence, the present mitogenome based phylogenies are congruent with the prevailing hypothesis and strengthen the evolutionary thoughts of Testudines in the Vertebrate tree of life.

**Fig 4.**
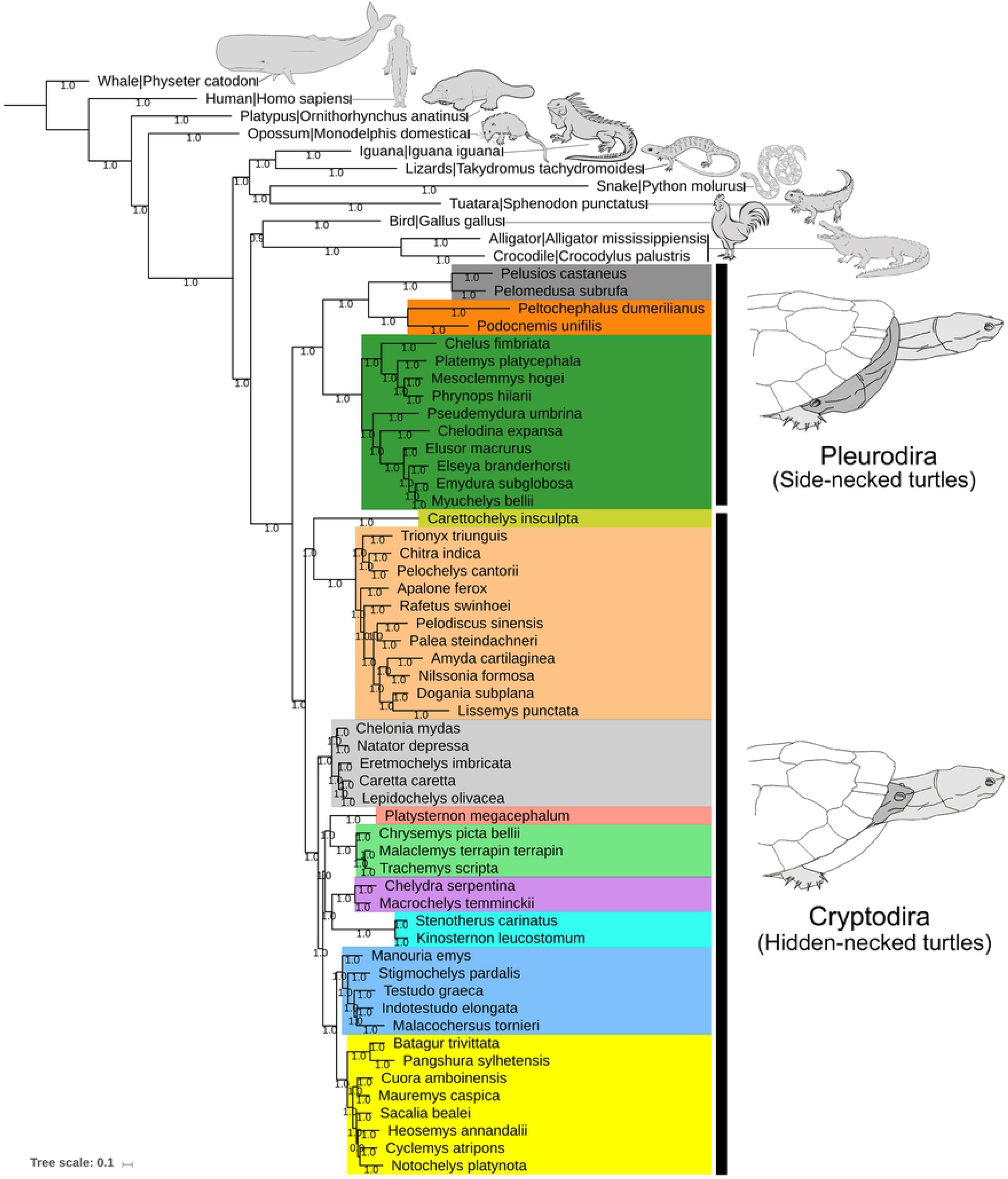
Bayesian (BA) phylogenetic tree based on the concatenated 13 PCGs of Testudines and other amniotes mitochondrial genomes indicated the diapsid affinities of Testudines and close relationship with Archosaurians. Color boxes indicate the family level clustering for the studied Testudines species and two sub-orders are marked by black bar. The BA tree is drawn to scale with posterior probability support values were indicated along with the branches. Representative organism line diagrams are acquired from cyberspace.

### Gene arrangements

To define the phylogenetic clustering at different taxonomic levels, and presume the evolutionary pathways among Testudines, TreeRex analysis was adopted to assure the gene arrangements. A total of 50 consistent nodes were detected in the present analysis (Fig. 5). Considering A50 node as an ancestral trait of both Cryptodiran and Pleurodiran species in the present dataset, the gene arrangement events are plesiomorphy for most of the Testudines species with few exceptions. Four gene arrangements events were detected in the present dataset: (i) an inversion of trnP was observed in A39 node of *E. macrurus*, (ii) an inversion of trnP was observed in A21 node of *E. imbricata*, (iii) two inversion of trnP and trnS1 were observed on node A11 towards A10 which separates the family Testudinidae from Geoemydidae, (iv) two inversion of trnS1, trnP and one TDRL event towards *P. megacephalum* separates the family Platysternidae to Emydidae (Fig. 5). However, the synapomorphy was observed in three species under three different families, Chelidae (*E. macrurus*), Cheloniidae (*E. imbricata*), and Platysternidae (*P. megacephalum*). As compared the gene arrangement scenario with other species under Chelidae and Cheloniidae, the GO of both *E. macrurus* and *E. imbricata* might occurr due to the parallel evolution, which needs further investigation. Further, to know the phylogenetic position and evolutionary history of the sole member (*P. megacephalum*) under the monotypic family Platysternidae, the mitogenomes data has been largely assessed. The unusual gene arrangements, duplication of CR, and loss of redundant genes was observed in *P. megacephalum* mitogenome [50,51,53]. Later on, the micro-evolutionary analysis was performed to know the evolution of CRs, and evidenced that, the duplicate CR in *P. megacephalum* derived from the heterological ancestral recombination of mitochondrial DNA [52]. The present TreeRex based GO analysis revealed that, both inversion and TDRL events plays a major role in the independent evolution of *P. megacephalum* as observed in the earlier studies. In the recent past, the draft genome of this species has been assembled to know the species even better [72]. Hence, we recommended whole genome data of Testudines and their comparative analysis will be essential to comprehend the molecular mechanisms leading to their distinctive morphology and physiology. Altogether, the structural features of mitochondrial genomes, phylogeny, and TreeRex based gene arrangement analysis fortify the knowledge on the Testudines evolution.

**Fig 5.**
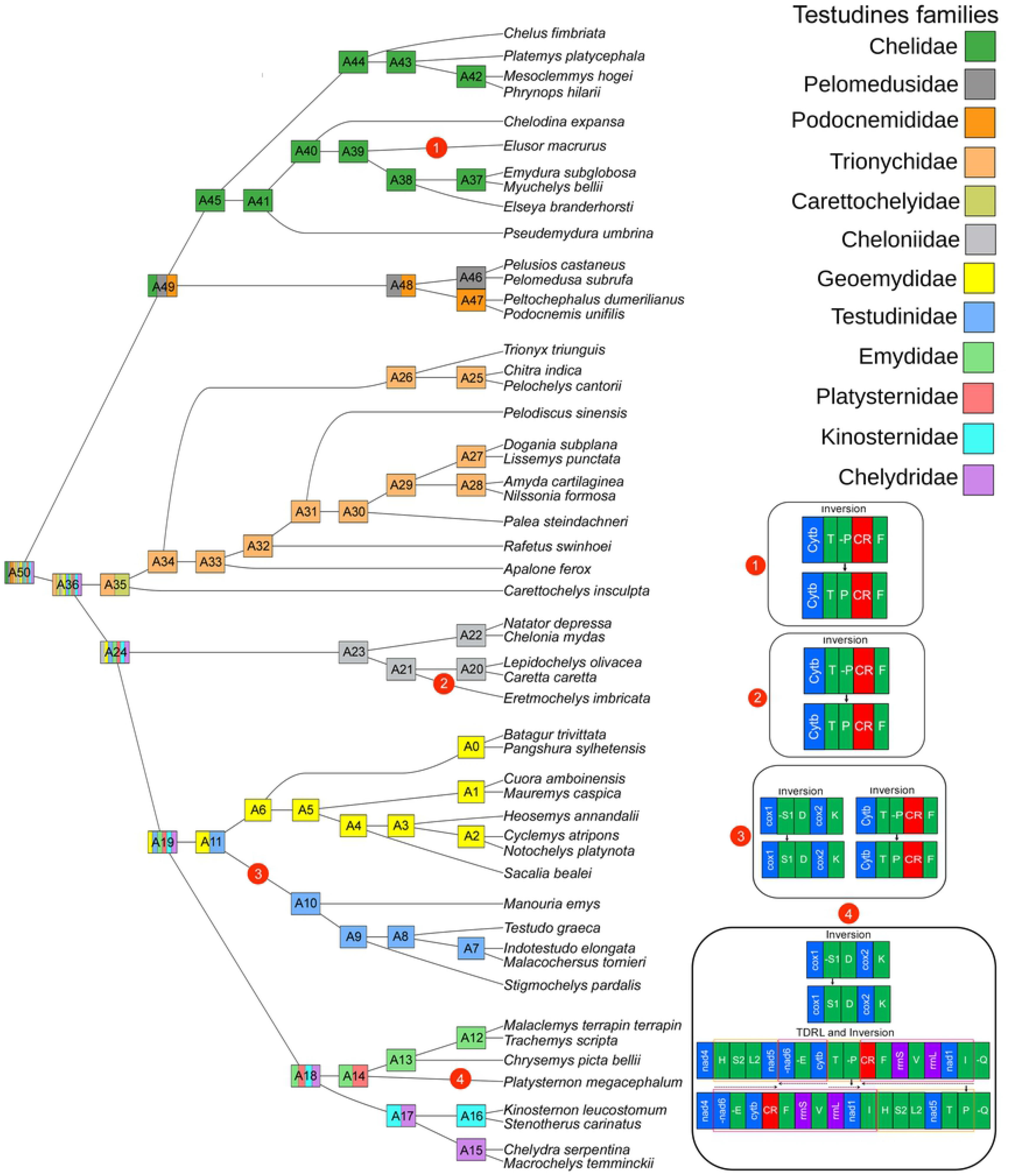
Gene-order based topology based on TreeREx analysis revealed the gene arrangement scenario within 52 Testudines species. All internal nodes are shared color as per phylogenetic clustering. Four gene arrangement scenarios are superimposed besides the tree.

### Conclusions and conservation inferences

The evolutionarily distinct and globally endangered (EDGE) freshwater turtle, *P. sylhetensis* (family Geoemydidae) is endemic to India and Bangladesh. Due to the habitat fragmentation, anthropogenic threats, and illegal poaching; the populations of *P. sylhetensis* have remarkably declining in its range distribution [73–75]. Hence, on priority, *P. sylhetensis* is categorized as an ‘endangered’ species in the IUCN Red data list, ‘Appendix II’ category in the Convention on International Trade in Endangered Species of Wild Fauna and Flora (CITES), and ‘Schedule I’ species in Indian Wildlife (Protection) Act, 1972. Nevertheless, owing to the unavailability of in-depth genetic information of many Testudines; species-specific biological information has not been amply resolved. The present study assembled and characterized the first complete mitogenome of *P. sylhetensis* (16,568 bp) and deliberated the comparative analysis with other 52 Testudines representing 12 families and two sub-orders (Cryptodira and Pleurodira). The estimated PCGs based BA and ML phylogenies indicated that, *P. sylhetensis* is closely related to *B. trivittata* and congruent with the earlier study [62]. Nevertheless, the phylogenetic reassessment with more mitogenomes, evidenced diapsid affinity and sister relationship of Testudines with Archosaurians, which would be fortify the Vertebrate tree of life. The GO analysis also revealed that, most of the species mitogenomes accommodate similar gene arrangements as observed in typical vertebrates with few exceptions (*E. macrurus*, *E. imbricata*, *P. megacephalum*, and shared nodes of Geoemydidae-Testudinidae). Hence, the generated mitogenomic information of *P. sylhetensis* would be useful for further phylogenetic reconciliation, to understand the evolutionary history, and adopt the precise conservation management.

## Acknowledgements

We thank the Director of Zoological Survey of India (ZSI), Ministry of Environment, Forests and Climate Change (MoEF&CC), Govt. of India for providing necessary permissions and facilities. We are thankful to the Arunachal Pradesh Biodiversity Board for providing necessary permissions and facilities. The first author (S.K) acknowledges the fellowship grant received from Council of Scientific and Industrial Research (CSIR) Senior Research Associateship (Scientists’ Pool Scheme) Pool No. 9072-A.

## Author Contributions

Conceptualization: SK, VK; Data curation: KT, SK; Formal analysis: SK; KT; Funding acquisition: KC, VK; Investigation: SK, VK; Methodology: SK, KT, VK; Project administration: KC, VK; Resources: KC; Software: SK, KT; Supervision: VK, KC; Validation: SK, VK; Visualization: SK, KT; Writing – original draft: SK, KT, VK; Writing – review & editing: SK, VK, KC.

## Data Availability Statement

The following information was supplied regarding the accessibility of DNA sequences: The complete mitogenome of *Pangshura sylhetensis* is deposited in GenBank of NCBI under accession number MK580979.

## Supporting information

**S1 Fig.** A pictorial overview of the methodologies used for sequencing and analysis of *P. sylhetensis* mitogenome and bioanalyzer profiles after sonication of enriched mitochondrial DNA sample and libraries.

**S2 Fig.** Putative secondary structures for 22 tRNA genes in mitochondrial genome of *P. sylhetensis*. The first structure shows the nucleotide positions and details of stem-loop of tRNAs. The tRNAs are represented by full names and IUPAC-IUB single letter amino acid codes. Different base pairings are marked by red, blue and green color bars respectively.

**S3 Fig.** Comparison of control region (CR) stem-loop structures of the origin of L-strand replication of 24 Testudines mitochondrial genomes.

**S4 Fig.** Comparison of control region (CR) stem-loop structures of the origin of L-strand replication of 26 Testudines mitochondrial genomes.

**S5 Fig.** Maximum Likelihood (ML) phylogenetic tree based on the concatenated nucleotide sequences of 13 PCGs of 52 Testudines species. Color boxes indicate the family level clustering for the studied species. The ML tree is drawn by IQ-Tree with bootstrap support values were indicated along with each node.

**S6 Fig.** Maximum Likelihood (ML) phylogenetic tree based on the concatenated 13 PCGs of Testudines and other amniotes mitochondrial genomes. Color boxes indicate the family level clustering for the studied Testudines species. The ML tree is drawn by IQ-Tree with bootstrap support values were indicated along with each node.

**S1 Table.** List of mitogenome sequences of Testudines and other amniotes species acquired from the NCBI database.

**S2 Table.** Estimated models by partitioning the 13 PCGs separately through PartitionFinder 2 for phylogenetic analysis of two datasets (52 mitogenomes of Testudines and 63 mitogenomes of Testudines + other amniotes).

**S3 Table.** Gene arrangements of the studied Testudines species used in the TreeREx analysis.

**S4 Table.** Nucleotide composition of 52 Testudines species mitochondrial genomes. The A+T biases of whole mitogenome, protein coding genes, tRNA, rRNA, and control regions were calculated by AT-skew = (A-T)/(A+T) and GC-skew= (G-C)/(G+C), respectively.

**S5 Table.** Frequency of start and stop codon distribution within the complete mitogenomes of 52 Testudines.

